# Rab46: a novel player in mast cell function

**DOI:** 10.1101/2023.03.13.532191

**Authors:** Lucia Pedicini, Jessica Smith, Sinisa Savic, Lynn McKeown

**Affiliations:** Leeds Institute of Cardiovascular and Metabolic Medicine, Faculty of Medicine and Health, University of Leeds, LS2 9JT, UK; Department of Clinical Immunology and Allergy, St James’s University Hospital, Leeds, United Kingdom; National Institute for Health Research-Leeds Biomedical Research Centre and Leeds Institute of Rheumatic and Musculoskeletal Medicine, Wellcome Trust Brenner Building, St James’s University Hospital, Leeds, United Kingdom

**Keywords:** mast cells, EFCAB4B, Rab46, CRACR2A, histamine, degranulation, trafficking, Rab GTPases

## Abstract

Mast cells are infamous for mediating allergic and inflammatory diseases due to their capacity of rapidly releasing a wide range of inflammatory mediators stored in cytoplasmic granules. However, mast cells also have several important physiological roles that involves selective and agonist-specific release of these active mediators. Whilst a filtering mechanism at the plasma membrane could regulate selective release of some cargo, the plethora of stored cargo and the diversity of mast cell functions suggests the existence of granule subtypes with distinct trafficking pathways. The molecular mechanisms underlying differential trafficking and exocytosis of these granules are not known, neither is it clear how granule trafficking is coupled to the stimulus. In endothelial cells, a Rab GTPase, Rab46, responds to histamine but not thrombin signals, and this regulates the trafficking of a subpopulation of endothelial-specific granules. Here, we sought to explore if Rab46 has a similar function in mast cells. We demonstrate, for the first time, that Rab46 is highly expressed in human and murine mast cells and Rab46 genetic deletion has an effect on mast cell degranulation that depends on both stimuli and mast cell developmental stage. Rab46 could therefore be an important regulator of stimuli-coupled responses in mast cells and future studies will seek to understand these mechanisms in order to develop novel and specific therapeutic targets for treatment of the diverse pathologies mediated by mast cells.

## Introduction

Mast cells store a powerful arsenal of pro-inflammatory mediators that, upon release from the cells, mediate a variety of physiological functions such as wound healing in response to injury and the development of acute inflammation [1]. Mast cells are also directly involved in several pathological conditions and are prominent initiators of allergic diseases [2]. Crosslinking of IgE on the primed mast cell surface causes the characteristic symptoms of allergy by the release of histamine, cytokines, leukotrienes, proteases and heparin from mast cell granules. However, in order to deliver unique physiological responses, it is becoming increasingly clear that selective release of granule cargo is distinctly coupled to the activating signal, the dysregulation of which leads to the pathological conditions associated with aberrant mast cell degranulation [3]. There are several studies exploring the effects of distinct agonists on the differential release of these pro-inflammatory cargo [4], however, little is known about the molecular mechanisms underlying the trafficking and secretion of the secretory granules (SGs) in which these mediators are stored [5]. An essential step toward deciphering the mechanisms behind exocytosis is the identification of the cellular components that regulate this process. Rab GTPases are master regulators of intracellular trafficking events and thereby they regulate immunity and inflammation responses by controlling granule trafficking and secretion in immune cells [6]. Several conventional Rab proteins are reported to be involved in mast cell exocytosis however, new evidence has recently pointed to the contribution of a new class of large EF-hand containing Rab GTPases in mast cells degranulation and function [7]. Rab46 (CRACR2A-L; *Ensembl* CRACR2A-203; CRACR2A-a) is a large Ca^2+^-sensing GTPase that, in addition to a Rab-domain, contains a coil-coiled domain and two EF-hand domains [8]. Rab46 is necessary for stimuli-coupled trafficking of selective cargo-restricted endothelial vesicles to the microtubule organising centre [9, 10] and for vesicle translocation to the immunological synapse in T-cells [11]. Here, we investigated the role of Rab46 in regulating mast cell degranulation and in establishing the pro-inflammatory environment typical of many immune-related diseases.

Rab46 (732 amino acids) is one of two functional isoforms translated by the EFCAB4B gene. CRACR2A (CRACR2A-S; *Ensembl* CRACR2a-201; CRACR2A-c), is a shorter (395 amino acids), non-Rab isoform of EFCAB4B expressed in T-cells and regulates store-operated calcium entry and T-cell signalling [12]. Analysis of EFCAB4B isoform expression in mast cells suggests Rab46 is expressed in both murine and human mast cells (Fig. 1a-b). Quantitative qPCR data suggest that Rab46 mRNA expression is five times more abundant than CRACR2A in both murine bone-marrow derived mast cells (BMMCs) and peritoneal derived mast cells (PMCs) (Fig. 1b). Moreover, and of particular interest because of the emerging role of these EF-hand containing Rab GTPases in mast cell function [7], we found that among the three known large Rab GTPases (Rab44, Rab45, Rab46), Rab46 mRNA is the most abundant. Rab46 mRNA is seven times more abundant than Rab44 in both BMMCs and PMCs, whereas Rab45 mRNA was not detected (Fig. 1c).

**Figure 1.**
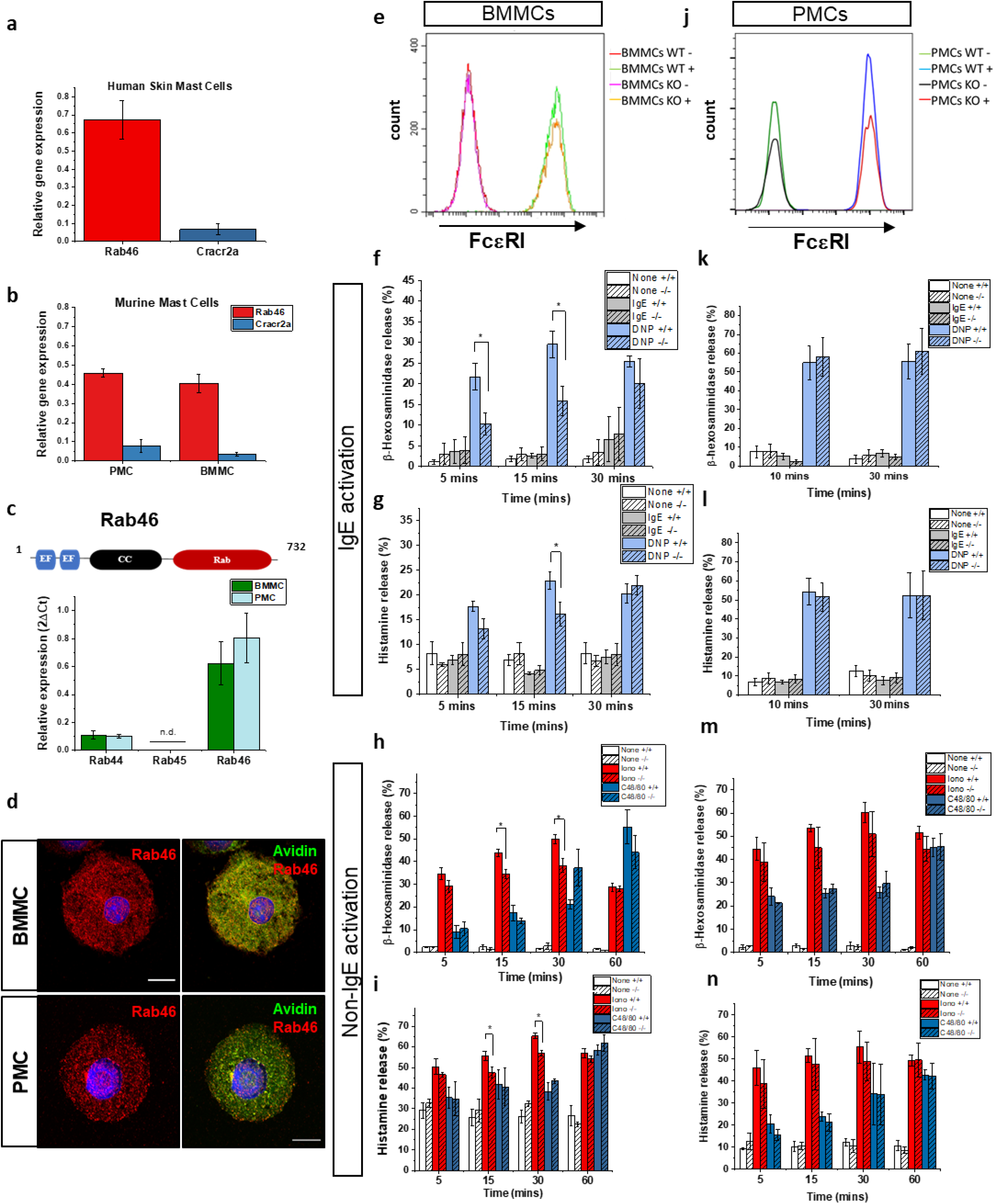
The role of Rab46 in mast cell degranulation. qPCR analysis of relative Rab46 mRNA expression in human mast cells (a) and murine peripheral mast cells (PMCs) and marrow mast cells (BMMCs) (b). (c) Schematic of Rab46 domains and relative mRNA expression of similar large Rab GTPases Rab44, Rab45 and Rab46. (d) Immunofluorescent imaging of Rab46 (red) and avidin (green) in PMCs and BMMCs. Nuclei = Hoescht (blue). (e) FACs analysis of FcεRI expression in BMMCs extracted from Rab46^wt^ and Rab46^-/-^ mice. (f-g) Time-dependent measurement of IgE-mediated β-hex (f) and histamine (g) release in BMMCs extracted from Rab46^wt^ and Rab46^-/-^ mice. Cells acutely pre-incubated with anti-DNP-IgE were treated with human serum albumin DNP (h-i) Time-dependent measurement of ionomycin and c48/80-mediated β-hex (f) and histamine (g) release in BMMCs extracted from Rab46^wt^ and Rab46^-/-^ mice. (j) FACs analysis of FcεRI expression in BMMCs extracted from Rab46^wt^ and Rab46^-/-^ mice. (k-l) Time-dependent measurement of IgE-mediated β-hex (k) and histamine (l) release in PMCs extracted from Rab46^wt^ and Rab46^-/-^ mice. (m-n) Time-dependent measurement of ionomycin and c48/80-mediated β-hex (m) and histamine (n) release in PMCs extracted from Rab46^wt^ and Rab46^-/-^ mice.

In our previous studies, we demonstrated the role of Rab46 in the trafficking of selective endothelial granules and demonstrated the colocalization of Rab46 with granule markers [9]. Here, we observed localization of Rab46 to mast cell granules (Fig. 1d). Confocal analysis of both BMMCs and PMCs revealed that endogenous Rab46 (in red) localizes to mast cell granules stained with avidin (in green). These results suggest a potential role of Rab46 in the regulation of mast cells trafficking.

To determine the effect of Rab46 depletion on mast cell degranulation we measured β-hexosaminidase (β-hex) and histamine secretion in BMMCs and PMCs extracted from wild-type (Rab46^wt^) mice and mice where Rab46 had been depleted (Rab46^-/-^), using IgE and non-IgE stimuli. Firstly, we measured mast cell secretion upon stimulation of the high-affinity IgE receptor (FCεRI) using acute IgE-sensitized BMMCs. FcεRI expression was comparable between BMMCs extracted from Rab46^wt^ and Rab46^-/-^ mice (Fig. 1e). However, in mast cells pre-treated for 1 hour with anti-DNP IgE, DNP-human serum albumin (HSA)-evoked β-hex and histamine secretion was significantly reduced in mast cells derived from Rab46^-/-^ mice compared to Rab46^wt^ (Fig. 1f-g). Measurement of non-immunological activation of mast cells by the calcium ionophore, ionomycin, also demonstrated that Rab46 depletion induces a significant reduction in both β-hex and histamine release. In addition to FcϵRI, mast cell degranulation occurs via activation of Mas-related G protein-coupled receptor X2 (MRGPRX2; mouse ortholog MrgprB2). Here, using the MRGPRX2 specific agonist compound 48/80 (C48/80) (Fig. 1h-i), Rab46 deletion had no significant effects on β-hex and histamine secretion in BMMCs. These results suggest a role for Rab46 in IgE (FcεRI)-mediated secretion and ionomycin-mediated mast cell degranulation but not the specific MRGPRX2 pathway in BMMCs.

Interestingly, although Rab46 expression in PMCs is comparable to BMMCs and PMCs exhibit a robust IgE-mediated response, there was no significant differences observed in IgE-mediated β-hex and histamine release between mast cells extracted from Rab46^wt^ and Rab46^-/-^ mice (Fig. 1k-l). In addition, no significant differences were detected in ionomycin stimulated β-hex and histamine release between PMCs extracted from Rab46^wt^ and Rab46^-/-^ mice (Fig. 1m-n).

These data suggest that PMCs exhibit a stronger and faster degranulation response compared to BMMCs, however, the response to the stated agonists appears to be independent of Rab46. On the other hand, Rab46 deficiency significantly impairs BMMC degranulation upon FcεRI and ionomycin stimulation while activation of the MRGPRX2 pathway triggers a Rab46-independent response in both BMMCs and PMCs.

In this study, we demonstrate for the first time the expression of Rab46, a novel EF-hand containing GTPase in mast cells and our in vitro results suggest a role for Rab46 in exocytosis in BMMCs. Interestingly, in endothelial cells, Rab46 is coupled to specific agonists to mediate the selective trafficking of sub-populations of endothelial-specific vesicles. Both endothelial-specific vesicles and mast cell granules are derived from lysosomes, indicating they may share common regulatory mechanisms. Here, the data implies that Rab46 may also play a role in agonist-specific pathways in mast cells. Furthermore, the data suggests Rab46 may regulate granule release in sub-populations of mast cells dependent on developmental stage or mast cell subtype, underlying the importance of mast cell heterogeneity. A greater understanding of Rab46-dependent trafficking events will provide opportunities to develop novel therapeutic targets for the treatment of the increasing prevalence of allergic and other inflammatory diseases.

## Acknowledgements

We thank Leeds Bioimaging and FACs facility staff for their support. We particularly would like to thank Prof. Marcus Maurer and Dr Joerg Scheffel, Charite - University of Berlin, for giving us time and training in their laboratory.

## Online Materials and Methods

### Ethics

Murine work was carried out in accordance with The Animals (Scientific Procedures) Act 1986 (Amended 2012) and all protocols were authorized by the University of Leeds Animal Ethics Committee and Home Office UK (Project License P606320FB to David J Beech). Mice were kept in the University of Leeds animal facility under standard conditions, including a 12-hour sleep/wake cycle, with access to water and chow diet ad libitum.

### Mice

Rab46^-/-^ cells are from CRACR2A tm1.1 (KOMP) vlcg (global knockout mice that were created from ES cell clone 15424A-C4, generated by Regeneron Pharmaceuticals, Inc. and obtained from the KOMP Repository (www.komp.org). Rab46^+/+^ wild-type cells are derived from C57BL/6N black controls. Genotypes of the mice were validated using real-time PCR with specific probes designed for each gene (Transnetyx, Cordova, TN).

### Cell culture

Bone marrow cells from 8-week-old female mice were cultured at 37°C and 5% CO_2_ in RPMI 1640 medium containing 10% fetal bovine serum, 2 mM L-glutamine, non-essential amino acids, 1% penicillin and streptomycin, and supplemented with 20 ng/ml interleukin-3 and 20 ng/ml Stem Cell Factor (both from Peprotech, Rocky Hill, NJ). After four weeks in culture, more than 95% of the cells were mast cells as assessed by FcεRI and c-kit expression by flow cytometry.

Peritoneal cells were obtained by injecting the mice with 5 ml of PBS for the peritoneal lavage. Cells were centrifuged at ∼300 × g and resuspended in RPMI Medium supplemented with 10% Fetal Calf Serum (FCS), 1% PenStrep, non-essential amino acids, 20 ng/ml IL-3, and 20 ng/ml SCF. Cells were further cultured in 5% CO_2_ at 37°C. On the 2nd day of cultivation, all non-adherent cells were collected and transferred into a new flask. PMCs were used for experiments between 10 and 25 days of culturing. Flow cytometry analysis identified 98.5–99.5% cells to be double positive for FcεRI and c-Kit and could be ranked as mast cells.

### RT-PCR

RNA isolation was performed using the High Pure RNA isolation kit (Roche) according to manufacturer’s protocol. Reverse transcription (Applied Biosystems) was followed by real-time PCR using SYBR green using a LightCycler (Roche). Expression levels of housekeeping genes (GAPDH and b-actin) were also measured. Primer sequences are provided in table below

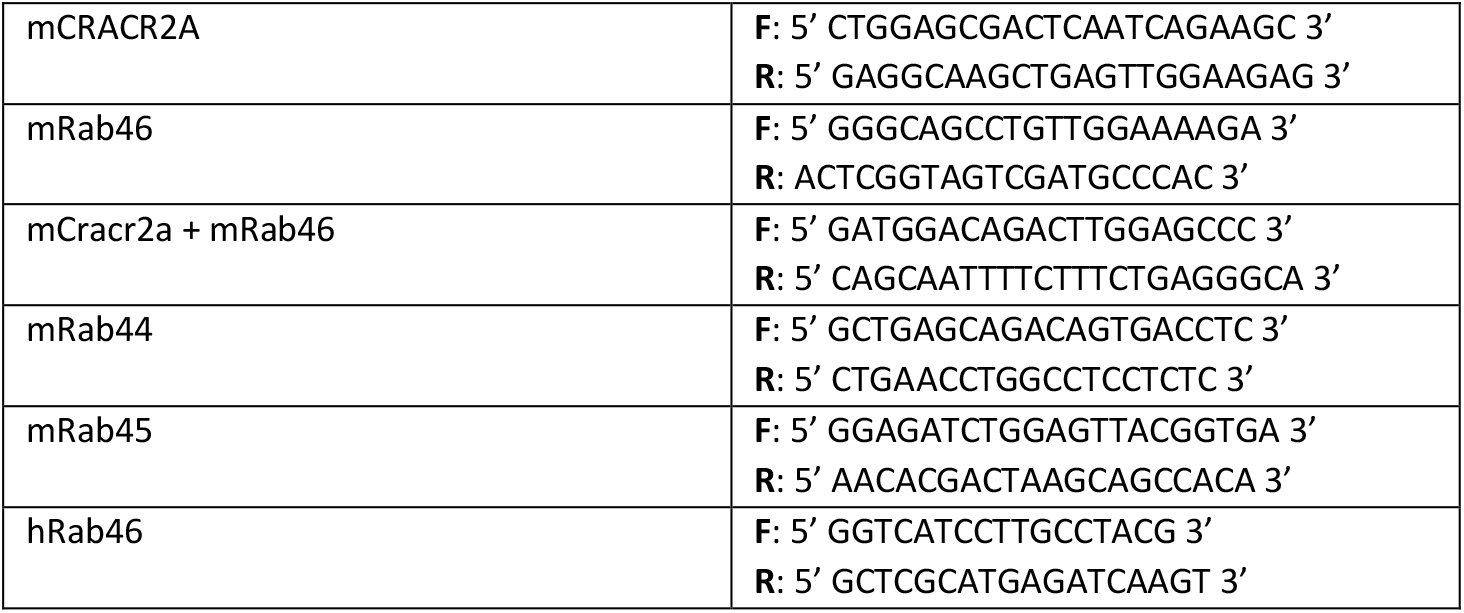

### Immunocytochemistry

A total of 5 × 10^4^ MCs were plated on Cell-Tak (Corning)-coated Ibidi 8-well plates. Cells were fixed with 4% PFA and secretory granules were stained with avidin-FITC (BioLegend) and rabbit anti-Rab46 (CRACR2A) antibody (ProteinTech) for 1 hour at RT. After washing in PBS, a fluorescently labelled secondary antibody (Alexa Fluor 594 anti-goat IgG - Jackson ImmunoResearch Labs) was applied for 30 mins. Cells, washed with PBS, were briefly incubated in Hoechst before being mounted with Ibidi mounting medium.

### High-resolution AiryScan microscopy

High-resolution microscopy was performed using an inverted confocal laser-scanning microscope, Zeiss LS880 with AiryScan system. Images were captured using a 63×/1.4 oil objective and 405-nm diode, Argon/2 (458, 477, 488, and 514 nm), HeNe 543 nm, and HeNe 633 nm lasers. All the images were acquired and processed with Zen software. All imaging was performed at room temperature.

### Flow cytometry

PMCs and BMMCs were centrifuged at 300 g for 5 mins and 1 × 10^5^ cells per reaction re-suspended in 200 µl FACs buffer (500 ml PBS; 2 % FBS; 1 mM EDTA; 25 mM HEPES; pH 7.4) in a FACS tube. Cells placed on ice and incubated with 1:100 PE-tagged anti-PE-FCεRIa (BioLegend) or APC-CD117 (BioLegend) or negative control for 20 mins. After 3 x wash, cells were resuspended in 300 µl FACs buffer and fluorescent measurements taken on a CytoFlex 2 laser flow cytometer. Analysis and graphs created using FlowJo software.

### Degranulation Assay

PMCs and BMMCs were centrifuged at 300 g for 5 mins and re-suspended in Tyrodes solution containing in mM: NaCl 135; KCl 5; CaCl_2_ 1.8; MgCl_2_ 1; glucose 5.6; HEPES 20; pH 7.4. The cells were seeded in a U-bottom 96-well plate (5 × 10^5^ cells/well). All experimental conditions were performed in triplicates. For sensitization, 1 × 10^6^ MCs were incubated with 1 μg/ml of monoclonal anti-DNP IgE (Sigma) for 1 h at 37°C. Degranulation was induced by incubation of BMMCs in the presence of 100 ng/ml Dinitrophenyl human serum albumin (DNP) (10 ng/ml PMCs) or 1 µg/ml IgE or 10 µM C48/80 or 2 µM ionomycin or buffer controls at 37°C and 5% CO_2_. Cells were centrifuged at 300 g for 5 min. The supernatants were separated and the cell pellets were lysed in ddH2O during 5 min at room temperature and then frozen at -80 C.

### Beta-hexosaminidase release

The amount of released β-hexosaminidase enzyme was quantified by spectrophotometric analysis of 4-Methylumbelliferyl N-acetyl-β-D-glucosaminide (4-MUG) hydrolysis, as previously described (ref). In short, the cell lysates and supernatants were incubated separately with 5 µM 4-MUG in 100mM citrate buffer for 1 h at 37°C and the reaction was stopped by adding 100 mM Na2CO3-NaHCO3 buffer (pH 10.7). Hydrolysis rate of 4-MUG was quantified by fluorometric measurements at 460 nm emission and 355nm excitation using the Synergy H1 spectrophotometer. The percentage β-hexosaminidase release was calculated as the absorbance ratio of the supernatant to the sum of supernatant and lysate. If not stated otherwise, all chemicals used in this work were purchased from Sigma.

### Quantitation of histamine

Histamine content in the supernatant and the pellet was measured by a fluorometric method based on its stable condensation with O-phthalaldehyde (OPT) in an alkaline medium. Briefly, supernatants and intracellular contents were collected and exposed to 2.3 μl OPT (7.5 μM) and 9 μl NaOH (1 mM) for 4 min. After stabilization with 3 µl of HCl (3 mM), fluorescence at 360 nm in a conventional spectrophotometer was quantified.

### Statistics

All average data are represented by mean ± SEM. Paired *t* test was performed as appropriate when comparison between two data groups was sought. Statistical significance was considered to exist at P < 0.05 (*, P < 0.05; **, P < 0.01). Where comparisons lack an asterisk, they were not significantly different and/or marked as not significant. OriginPro 2017 software was used for data analysis and presentation.

## References

1. Dahlin, J.S., et al., The ingenious mast cell: Contemporary insights into mast cell behavior and function. Allergy, 2022. 77(1): p. 83–99.

2. Marquardt, D.L. and S.I. Wasserman, Mast cells in allergic diseases and mastocytosis. West J Med, 1982. 137(3): p. 195–212.

3. Gaudenzio, N., et al., Different activation signals induce distinct mast cell degranulation strategies. J Clin Invest, 2016. 126(10): p. 3981–3998.

4. Yu, Y., et al., Non-IgE mediated mast cell activation. Eur J Pharmacol, 2016. 778: p. 33–43.

5. Moon, T.C., A.D. Befus, and M. Kulka, NMast cell mediators: their differential release and the secretory pathways involved. Front Immunol, 2014. 5: p. 569.

6. Prashar, A., et al., Rab GTPases in Immunity and Inflammation. Front Cell Infect Microbiol, 2017. 7: p. 435.

7. Kadowaki, T., et al., The large GTPase Rab44 regulates granule exocytosis in mast cells and IgE-mediated anaphylaxis. Cell Mol Immunol, 2020. 17(12): p. 1287–1289.

8. Wilson, L.A., et al., Expression of a long variant of CRACR2A that belongs to the Rab GTPase protein family in endothelial cells. Biochem Biophys Res Commun, 2015. 456(1): p. 398–402.

9. Miteva, K.T., et al., Rab46 integrates Ca(2+) and histamine signaling to regulate selective cargo release from Weibel-Palade bodies. J Cell Biol, 2019. 218(7): p. 2232–2246.

10. Pedicini, L., et al., Affinity-based proteomics reveals novel binding partners for Rab46 in endothelial cells. Sci Rep, 2021. 11(1): p. 4054.

11. Woo, J.S., et al., CRACR2A-Mediated TCR Signaling Promotes Local Effector Th1 and Th17 Responses. J Immunol, 2018. 201(4): p. 1174–1185.

12. Srikanth, S., et al., A novel EF-hand protein, CRACR2A, is a cytosolic Ca2+ sensor that stabilizes CRAC channels in T cells. Nat Cell Biol, 2010. 12(5): p. 436–46.

